# Reprogramming Dedifferentiation Regulatory Networks Preserves Human Chondrocyte Phenotypes

**DOI:** 10.64898/2026.05.10.724134

**Authors:** Ellen Y. Zhang, Sang Hyun Lee, Yu-Chung Liu, Yuna Heo, Hannah H. Kim, Dong Hwa Kim, Tyler E. Blanch, Jaeun Jung, Claudia Loebel, Melike Lakadamyali, Robert L. Mauck, Inkyung Jung, Su Chin Heo

**Affiliations:** McKay Orthopaedic Research Laboratory, Department of Orthopaedic Surgery, Perelman School of Medicine, University of Pennsylvania, Philadelphia, PA, 19104, USA; Department of Bioengineering, School of Engineering and Applied Science, University of Pennsylvania, Philadelphia, PA, 19104, USA; Department of Physiology, Perelman School of Medicine, University of Pennsylvania, Philadelphia, PA, 19104, USA; Epigenetics Institute, University of Pennsylvania, Philadelphia, PA, 19104, USA; Biochemistry and Molecular Biophysics Graduate Group, Perelman School of Medicine, University of Pennsylvania, Philadelphia, PA, 19104, USA; Department of Biological Sciences, Korea Advanced Institute of Science and Technology (KAIST), Daejeon, 34141, KOR

**Keywords:** chondrocyte dedifferentiation, autologous chondrocyte implantation, chromatin remodeling, single-nucleus multiome, cartilage regeneration

## Abstract

Chondrocyte-based cartilage repair strategies such as autologous chondrocyte implantation (ACI) require extensive in vitro expansion to obtain clinically relevant cell numbers. However, this expansion step progressively drives chondrocyte dedifferentiation, reducing matrix-forming capacity and contributing to variable repair outcomes. To better understand this process, we used single-nucleus multiome profiling (snRNA-Seq + snATAC-Seq) to define the transcriptional and chromatin accessibility programs underlying human chondrocyte dedifferentiation during expansion. Multiome integration across passages revealed a continuous dedifferentiation trajectory accompanied by coordinated remodeling of gene expression and chromatin accessibility, identifying chromatin destabilization as an early regulatory event during phenotype loss. Guided by these regulatory signatures, we screened available small-molecule inhibitors targeting candidate pathways and found that Fludarabine most consistently preserved chondrocyte identity during early expansion. Fludarabine was associated with suppression of STAT1-related programs and early stabilization of the chromatin landscape prior to broader transcriptional recovery. Functionally, treated cells demonstrated enhanced matrix-forming capacity in chondrogenic pellet culture and significantly increased nascent protein synthesis in 3D hydrogel culture, with biosynthetic output approaching unexpanded controls by day 21. Together, these findings identify chromatin stability as a key regulatory determinant of expansion-associated chondrocyte dedifferentiation and establish a pharmacologic strategy to preserve chondrocyte functional potency during cell manufacturing for cartilage repair.

## 1. Introduction

Articular cartilage enables low-friction joint motion while sustaining and transmitting loads to the underlying subchondral bone over decades^1^. However, cartilage has a limited intrinsic capacity for repair, and injury or degeneration frequently progresses toward osteoarthritis (OA), a leading cause of chronic pain and impaired mobility^2^. The societal burden of cartilage degeneration is substantial, with arthritis-related costs exceeding hundreds of billions of dollars annually in the United States, and osteoarthritis prevalence continuing to rise^3^. Because mature cartilage is avascular and exists in a demanding mechanical environment, even focal defects often fail to heal effectively and can progress to joint-wide disease^2^.

For localized cartilage lesions, autologous chondrocyte implantation (ACI) is an established biological repair strategy^4^. In ACI and its more recent matrix-assisted variant (MACI), chondrocytes are harvested arthroscopically from a healthy, non–weight-bearing region and expanded *in vitro* for approximately three to five weeks before re-implantation into the defect^5^. ACI often demonstrates superior clinical outcomes compared with other approaches, such as microfracture, but its long-term durability remains variable^6^. A fundamental barrier is the need for substantial chondrocyte expansion. Because of the limited availability of primary tissue and the low cellularity of adult articular cartilage, ACI typically requires four to six population doublings *in vitro*. Although necessary, this expansion creates a core biological tradeoff: cells proliferate, but progressively lose their functional cartilage-forming identity.^7^.

This phenotype loss is commonly termed chondrocyte dedifferentiation, a process broadly characterized as the reverse of chondrogenic differentiation during *in vitro* expansion on tissue culture plastic^7^. Dedifferentiating chondrocytes undergo a phenotypic shift from a polygonal-shaped morphology toward a spindle-shaped, fibroblast-like appearance and downregulate canonical chondrogenic markers while showing reduced matrix production capacity^7–9^. These changes are accompanied by cytoskeletal remodeling and increased focal adhesion signaling, which have been implicated in the progression of dedifferentiation^10–12^. Functionally, these changes contribute to inferior repair tissue quality, with implanted cells more prone to forming mechanically weaker fibrocartilage rather than durable hyaline cartilage. Importantly, this phenomenon is not new, as dedifferentiation has been recognized for decades^7,8^, yet effective and clinically practical strategies to prevent it remain limited, and the mechanisms driving this phenotypic shift are still incompletely understood. Multiple interventions have been explored to preserve the chondrogenic phenotype during *in vitro* expansion, including manipulation of growth factors, culture conditions, and substrate properties^8,13–19^. While many of these approaches can partially mitigate dedifferentiation or promote redifferentiation, none have fully prevented dedifferentiation during extended expansion, limiting the ability to generate larger numbers of high-quality therapeutic chondrocytes. This unmet need underscores the importance of expansion-compatible interventions that preserve chondrocyte identity without compromising proliferative yield.

Beyond transcriptional marker changes, upstream regulatory remodeling at the epigenetic level may influence cell phenotype, including the stability and reversibility of phenotype loss^20,21^. In support of this concept, multi-omic approaches are increasingly being used to resolve continuous biological processes, including dedifferentiation trajectories, identify transient intermediate states, and map candidate regulators and pathways that drive phenotypic shifts from native to dedifferentiated states^22,23^. Critically, single-nucleus profiling is particularly well-suited for this context, as it captures cellular heterogeneity and resolves distinct transcriptional and chromatin states within a population that bulk analyses would obscure^22^.

In this study, we used single-nucleus multiome sequencing (snRNA-Seq + snATAC-Seq) to define the transcriptional and chromatin accessibility programs underlying chondrocyte dedifferentiation during *in vitro* expansion. We found that dedifferentiation proceeds as a continuous transition accompanied by coordinated gene expression changes and early chromatin remodeling, identifying transition-associated transcription factors (TFs) as candidate regulators of phenotype loss. Guided by these regulatory signatures, we then evaluated a pharmacologic approach to improving expansion-associated phenotype stability, focusing on inhibition of signal transducer and activator of transcription 1 (STAT1) using Fludarabine. Fludarabine treatment was associated with stabilization of the chromatin landscape, preservation of chondrocyte morphology, a shift of cartilage-associated transcriptional programs toward a more native-like state, and improved matrix-forming capacity in both pellet and 3D hydrogel culture systems, with population-wide recovery of nascent protein synthesis approaching unexpanded controls.

Here, by integrating morphological, chromatin, transcriptional, and matrix-level readouts in expanded chondrocytes, we aim to establish a practical and scalable strategy to improve the quality of expanded cells for cartilage repair applications. Ultimately, improving the quality of expanded chondrocytes could enhance ACI outcomes and broaden the therapeutic window for cell-based cartilage regeneration.

## 2. Results

### *In vitro* expansion drives coordinated transcriptional and chromatin remodeling during chondrocyte dedifferentiation

ACI requires extensive *in vitro* expansion of chondrocytes, a process that is well known to induce dedifferentiation and loss of cartilage phenotype, thereby limiting the therapeutic efficacy of cell-based cartilage repair^5,7–9,24^. Consistent with this long-recognized challenge, bulk RNA-seq analysis revealed extensive transcriptional remodeling during *in vitro* expansion (**Supplemental Fig. S1A)**, accompanied by a marked reduction in canonical cartilage markers (**Supplemental Fig. S1D, F**). Yet, bulk transcriptomic profiling cannot resolve the cellular heterogeneity or the underlying chromatin regulatory programs driving this phenotypic transition. To define the transcriptional and epigenomic regulatory landscape of chondrocyte dedifferentiation at single-cell resolution, we therefore performed single-nuclei multiome (snMultiome) profiling across the *in vitro* expansion process. To define factors underlying this process, primary human articular chondrocytes from four adult donors (41-55 years) were expanded from passage 0 (P0) to passage 6 (P6), and single-nucleus multiome (snMultiome) profiling (snRNA-Seq + snATAC-Seq) was performed at P0, P3, and P6 to capture coupled transcriptional and chromatin accessibility changes during expansion-associated phenotype loss (**Fig. 1A**). Integration of snRNA-Seq and snATAC-Seq datasets resolved three passage-associated chondrocyte states: native P0 chondrocytes, transitioning P3/P6 chondrocytes, and fully dedifferentiated P6 chondrocytes (**Fig. 1B**). While P0 nuclei formed a distinct cluster, P3 and P6 nuclei showed substantial overlap, indicating a shared dedifferentiated state and suggesting that major dedifferentiation-associated changes had already occurred by P3. Consistent with this, bulk RNA-seq analysis revealed few differentially expressed genes between P3 and P6 compared to the pronounced transcriptional shift observed between P0 and P3 (**Supplemental Fig. S1B, C**). These findings indicate that the major transcriptional remodeling associated with dedifferentiation is largely complete by P3. Notably, integrated bimodal analysis nevertheless still enabled separation of P3 and P6 states; while P3 and P6 nuclei showed substantial overlap when visualized by original passage identity (**Supplemental Fig. S2A**), clustering analysis identified a distinct P6-enriched population that was not captured by transcriptional data alone (**Supplemental Fig. S2B**).

**Figure 1.**
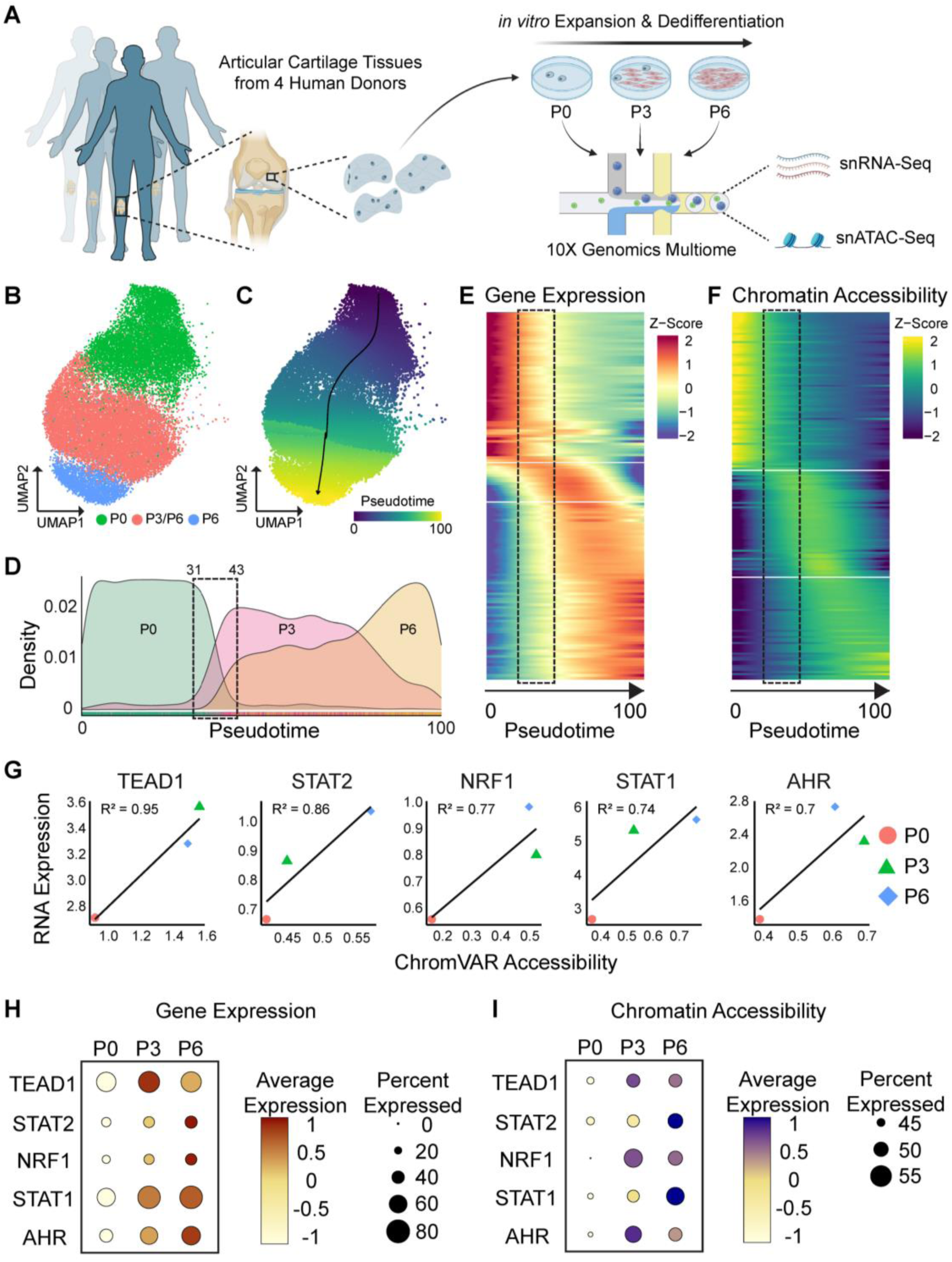
Integrated snMultiome profiling defines a dedifferentiation trajectory and identifies transcriptional regulators associated with chondrocyte expansion. **(A)** Schematic of the experimental design. Human articular chondrocytes isolated from 4 donors were expanded *in vitro* to P0, P3, and P6, followed by snMultiome profiling (paired snRNA-Seq and snATAC-Seq). **(B)** UMAP visualization of the integrated snMultiome dataset, colored by passage group (P0, P3, P6), and **(C)** pseudotime projection, showing the inferred dedifferentiation trajectory across *in vitro* expansion. The black line indicates the inferred trajectory path. **(D)** Density distribution of pseudotime values for P0, P3, and P6 nuclei, showing progressive shifts along pseudotime with passaging. Dashed box indicates the transitional window (pseudotime 31–43) capturing the phenotypic shift from the native to the dedifferentiated state. **(E)** Heatmap of pseudotime-ordered DEGs showing dynamic changes in gene expression across dedifferentiation. Dashed box highlights genes with dynamic expression changes within the transitional window (pseudotime 31–43). **(F)** Heatmap of pseudotime-ordered DARs showing dynamic changes in chromatin accessibility across dedifferentiation. Dashed box highlights regions with dynamic accessibility changes within the transitional window (pseudotime 31–43). **(G)** Correlation of TF RNA expression and chromatin accessibility (ChromVAR motif accessibility score) for representative candidate regulators (TEAD1, STAT2, NRF1, STAT1, and AHR) across passage groups. Linear regression fits and corresponding R^2^ values are shown. **(H)** Dot plot of RNA expression and (I) chromatin accessibility for selected TFs across P0, P3, and P6 nuclei. Dot color indicates average expression and dot size indicates the percentage of nuclei with detectable expression.

To model chondrocyte dedifferentiation as a continuous process rather than a discrete phenotype transition, we reconstructed a trajectory and assigned each nucleus a pseudotime value (**Fig. 1C**). This analysis identified a dominant trajectory spanning the native chondrocyte state to the late-expansion state, consistent with progressive shifts from P0 to P3 to P6 **(Fig. 1B, C)**. We next examined molecular features involved in the transition from the native P0 phenotype to the dedifferentiated P3/P6 state along the inferred trajectory. Nuclei were ordered by pseudotime, and pseudotime 31 to 43 was defined as a “transitional window” capturing the phenotypic shift (**Fig. 1D**). Visualization of pseudotime-ordered differentially expressed genes (DEGs) and differentially accessible regions (DARs) revealed coordinated changes in gene expression and chromatin accessibility across this transition (**Fig. 1E, F**).

To identify candidate transcriptional regulators associated with this progressive remodeling of regulatory programs during dedifferentiation, we integrated chromatin-based transcription factor (TF) activity (ChromVAR motif accessibility scores) with matched TF gene expression and evaluated their concordance across passage groups. Using this multimodal approach, we prioritized TFs that showed high expression and strong positive correlation between RNA expression and motif accessibility, including TEAD1, STAT2, NRF1, STAT1, and AHR, among others (**Fig. 1G; Supplemental Fig. S3**). TFs with high R^2^ values and a positive slope demonstrate coordinated increases in both transcriptional activity and chromatin accessibility across *in vitro* expansion, indicating progressive and concordant activation of these regulatory programs during dedifferentiation. Consistent with these findings, both RNA expression and chromatin accessibility for these TFs increased progressively from P0 to P6 (**Fig. 1H, I**). Together, these findings identify a set of candidate TFs whose activity is coordinately upregulated during expansion-associated chondrocyte dedifferentiation, providing regulatory nodes that may be targeted to preserve the chondrocyte phenotype.

### Fludarabine-mediated STAT1 inhibition mitigates chondrocyte dedifferentiation during *in vitro* expansion

Based on the candidate TFs identified in Figure 1, we next tested whether pharmacological inhibition of selected regulators could mitigate expansion-associated chondrocyte dedifferentiation (**Fig. 2A**). Small-molecule inhibitors were available for several candidate TFs, and among these, inhibition of STAT1 using Fludarabine produced the most consistent preservation of chondrogenic gene expression in the initial screen. STAT1 inhibition was therefore prioritized for downstream characterization. Results for all inhibitors tested, including treatment conditions and observations, are summarized in Supplemental Table S1 and Supplemental Fig. S4 (**Supplemental Table. S1, Supplemental Fig. S4**).

**Figure 2.**
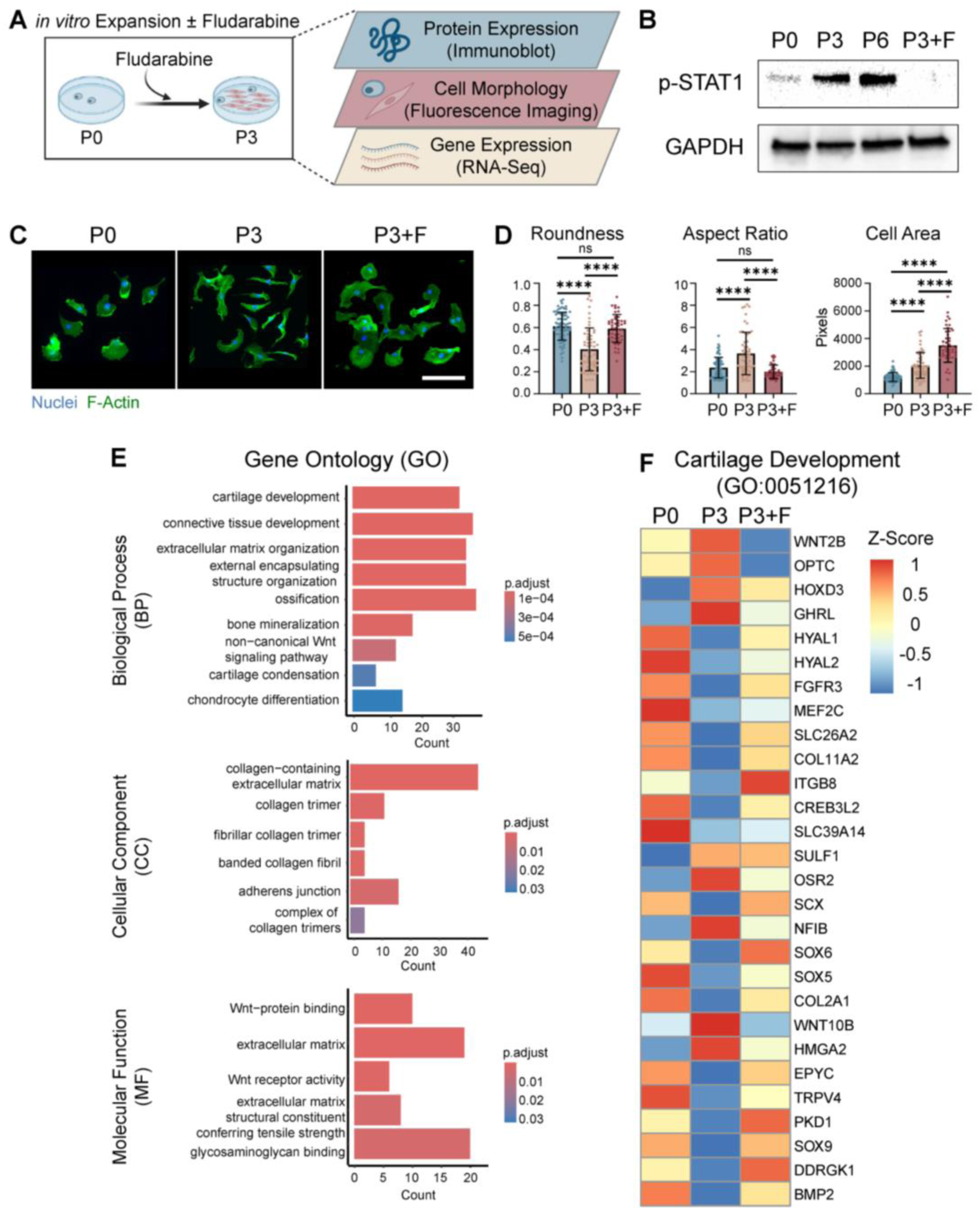
Fludarabine-mediated STAT1 inhibition mitigates chondrocyte dedifferentiation during early *in vitro* expansion. **(A)** Schematic of the Fludarabine treatment regimen and downstream assays used to evaluate chondrocyte preservation during *in vitro* expansion, including immunoblot, fluorescence imaging, and RNA-Seq. **(B)** Immunoblot of p-STAT1 and GAPDH in P0, P3, P6, and P3+F chondrocytes. **(C)** Representative fluorescence images of P0, P3, and P3+F chondrocytes stained for nuclei and F-actin. Scale bar = 100 µm. **(D)** Quantification of cell morphology metrics, including roundness, aspect ratio, and cell area, for P0, P3, and P3+F chondrocytes. Data presented as mean ± SEM from three donors; n = 75 for P0, n = 55 for P3, n = 46 for P3+F, ****p < 0.0001, ns = not significant (one-way ANOVA with Tukey’s post hoc test). **(E)** GO enrichment analysis of Fludarabine-rescued genes, showing enriched terms in biological process (BP), cellular component (CC), and molecular function (MF) categories. **(F)** Heatmap of genes in the GO term "cartilage development" (GO:0051216), showing expression shifts toward P0-like levels in P3+F cells relative to untreated P3 cells from three donors.

Because our snMultiome analysis indicated that major dedifferentiation-associated changes were already established by P3 (**Fig. 1B-D**), we used P3 as the primary endpoint for evaluating Fludarabine’s impact on chondrocyte preservation. To validate STAT1 engagement during dedifferentiation and to confirm effective inhibition of STAT1 by Fludarabine, we assessed phosphorylated STAT1 (p-STAT1) by immunoblotting. p-STAT1 levels increased during *in vitro* expansion and were reduced in Fludarabine-treated P3 cells (P3+F) relative to untreated P3 controls (P3) (**Fig. 2B**).

We next evaluated whether Fludarabine altered the expansion-associated morphological changes observed in monolayer culture. Consistent with previous reports^7–9,17,19^, untreated chondrocytes transitioned from a polygonal morphology at P0 to a more elongated, spindle-like morphology by P3 (**Fig. 2C, D**). Fludarabine treatment preserved a P0-like morphology, as shown by increased roundness and reduced aspect ratio relative to untreated P3 cells, although cell area remained elevated compared with P0 (**Fig. 2C, D**). To determine whether Fludarabine treatment affected cell proliferation during expansion, we assessed metabolic activity as a proxy for proliferation using a CCK-8 assay. Fludarabine-treated P3 chondrocytes (P3+F) showed proliferation comparable to untreated P3 controls, indicating that the phenotype-preserving effects of Fludarabine were not attributable to reduced proliferative activity (**Supplemental Fig. S4D**).

Next, to define how Fludarabine affects the transcriptome of chondrocytes, we performed bulk RNA sequencing (RNA-Seq) on P0, P3, and P3+F chondrocytes. Gene Ontology (GO) enrichment of biological process (BP), cellular component (CC), and molecular function (MF) categories identified recovery of terms related to chondrocyte phenotype, including cartilage development, chondrocyte differentiation, and extracellular matrix-associated pathways (**Fig. 2E**). Consistent with these pathway-level changes, chondrogenic genes within the GO term “cartilage development” (GO:0051216) showed broad recovery toward P0-like expression in P3+F cells (**Fig. 2F**). To determine whether Fludarabine could rescue chondrocytes that had already undergone dedifferentiation, we applied the inhibitor directly to P6 chondrocytes without prior continuous treatment. Late-stage Fludarabine addition did not restore a polygonal morphology or recover chondrogenic gene expression (**Supplemental Fig. S6**). These findings indicate that Fludarabine acts as a preventative agent rather than a rescue treatment.

Together, these data indicate that pharmacological inhibition of STAT1 using Fludarabine substantially preserves the chondrocyte phenotype during *in vitro* expansion, with coordinated effects on STAT1 activation, cell morphology, and gene expression.

### Fludarabine enhances functional cartilage matrix-forming capacity of expanded chondrocytes

Having established that Fludarabine-mediated STAT1 inhibition preserves chondrogenic gene expression, we next asked whether these transcriptional changes translated to functional preservation of cartilaginous matrix-forming capacity after prolonged *in vitro* expansion. To address this, primary human chondrocytes were expanded to P6 with (P6+F) or without (P6) Fludarabine treatment, formed into chondrocyte spheroids, and cultured for 7 days in chondrogenic medium with (CM+) or without (CM-) TGF-β3, a well-known potent growth factor that promotes chondrogenesis^25–27^ (**Fig. 3A**).

**Figure 3.**
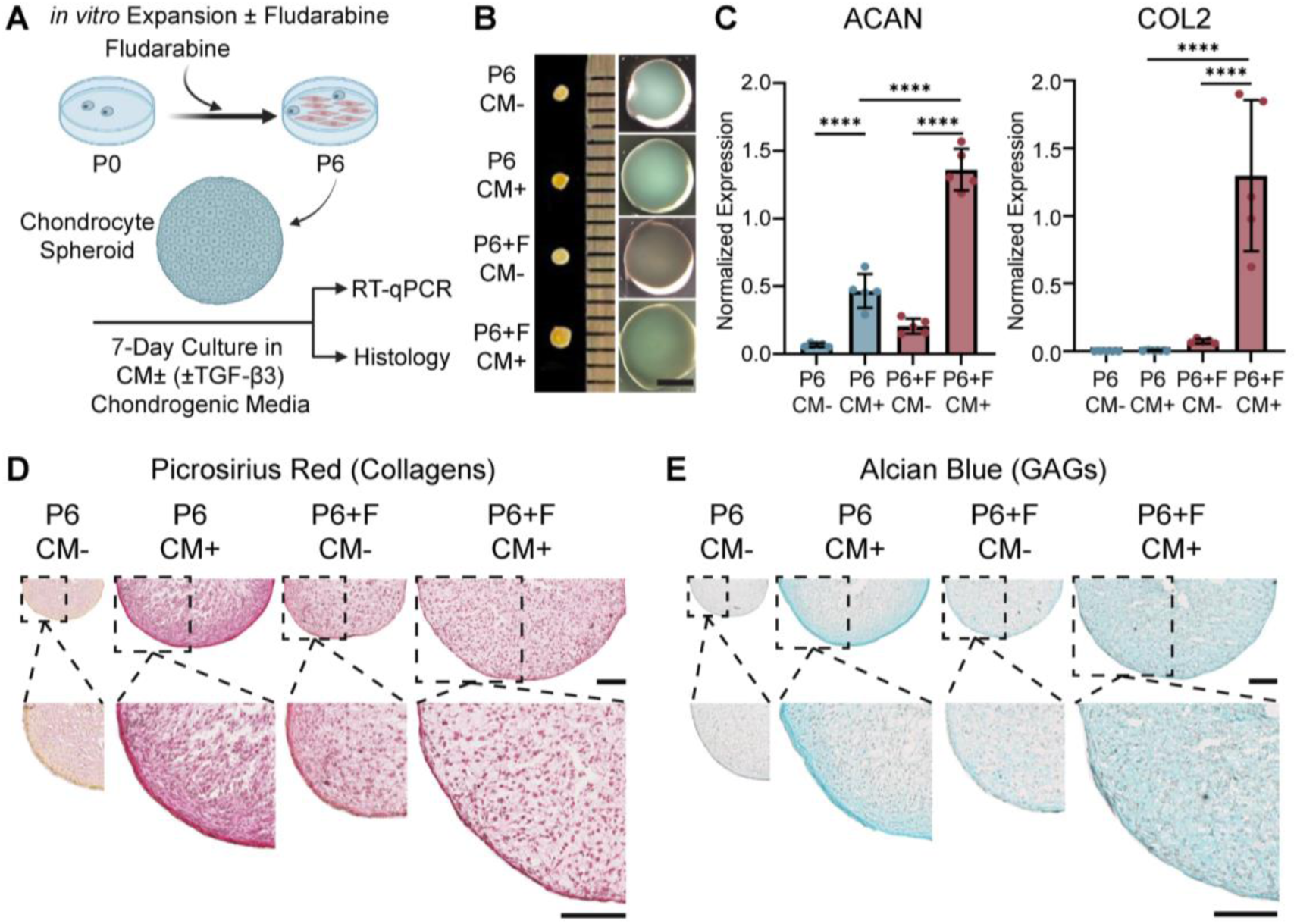
Fludarabine treatment during expansion improves chondrogenic redifferentiation and matrix formation in P6 chondrocyte spheroids. **(A)** Schematic of the experimental design. Human chondrocytes were expanded *in vitro* to P6 with or without Fludarabine treatment, formed into spheroids, and cultured for additional 7 days in chondrogenic medium with (CM+) or without (CM-) TGF-β3, followed by RT-qPCR and histological analysis. **(B)** Representative gross images of spheroids after 7 days of culture. Scale bar = 1 mm. **(C)** RT-qPCR analysis of ACAN and COL2 expression in P6 and Fludarabine-treated (P6+F) spheroids cultured under CM- or CM+ conditions. Data presented as mean ± SEM from one donor; n = 5, ****p < 0.0001 (one-way ANOVA with Tukey’s post hoc test). **(D–E)** Representative **(D)** PSR and **(E)** AB staining showing collagen deposition and glycosaminoglycan in spheroid sections. Dashed boxes indicate regions shown at higher magnification below. Scale bars = 250 µm.

Gross spheroid morphology at day 7 revealed clear differences in spheroid size and appearance among groups, indicating that both expansion history and chondrogenic stimulation influenced matrix-forming capacity (**Fig. 3B**). Fludarabine-treated expanded chondrocytes (P6+F) showed increased expression of chondrogenic genes, including aggrecan (ACAN) and type II collagen (COL2), even in the absence of TGF-β3 (CM-) (**Fig. 3C**). TGF-β3 stimulation (CM+) further enhanced chondrogenic gene expression in P6+F spheroids relative to untreated P6 controls, with the strongest induction observed in the P6+F CM+ group (**Fig. 3C**).

Histological analysis further supported enhanced matrix formation in Fludarabine-treated groups. Picrosirius red (PSR) staining revealed limited collagen deposition in untreated P6 spheroids cultured without TGF-β3 (P6 CM-), whereas the strongest and most uniform staining observed in P6+F CM+ spheroids (**Fig. 3D**). Similarly, Alcian blue (AB) staining demonstrated minimal glycosaminoglycan (GAG) deposition in P6 CM- spheroids, increased staining in P6 CM+, and further enhancement in P6+F groups, particularly in P6+F CM+ spheroids, which displayed the most robust and homogeneous matrix staining (**Fig. 3E**). In addition to increased staining intensity, Fludarabine-treated chondrocyte (P6+F CM+) spheroids displayed more uniform matrix distribution throughout the spheroid, consistent with improved cartilage-like matrix organization. In contrast, untreated P6 (P6 CM+) spheroids exhibited predominantly peripheral and heterogeneous matrix deposition (**Fig. 3D, E**).

Together, these findings demonstrate that Fludarabine-mediated STAT1 inhibition enhances the functional matrix-forming capacity of expanded chondrocytes and promotes more robust and uniform cartilage-like matrix organization.

### Fludarabine restores single-cell biosynthetic activity and cartilage matrix deposition in 3D culture

To further determine whether Fludarabine treatment improves matrix-forming function at the single-cell level after prolonged expansion, we next examined nascent protein synthesis and cartilage matrix marker deposition in NorHA hydrogel-encapsulated chondrocytes using fluorescent noncanonical amino acid tagging (FUNCAT) and immunofluorescence imaging^28,29^. Because 3D hydrogel systems provide a cartilage-mimetic microenvironment and are widely used platforms for cartilage regeneration^28,30,31^, this approach allowed us to assess whether Fludarabine-treated expanded chondrocytes retain functional biosynthetic activity in a physiologically relevant 3D context. To this end, chondrocytes expanded to P6 with or without Fludarabine were encapsulated in 5 kPa NorHA hydrogels and cultured for 21 days in CM+ medium supplemented with the noncanonical amino acid L-azidohomoalanine (AHA), with imaging performed at days 7 and 21 (**Fig. 4A**). AHA, a methionine analog, is incorporated into newly synthesized proteins in place of methionine, enabling nascent protein labeling^28,29^. Nascent proteins were detected by click chemistry using Dibenzocyclooctyne-amine-488 (DBCO-488) and cartilage matrix markers were detected by immunofluorescence staining (**Fig. 4B**).

**Figure 4.**
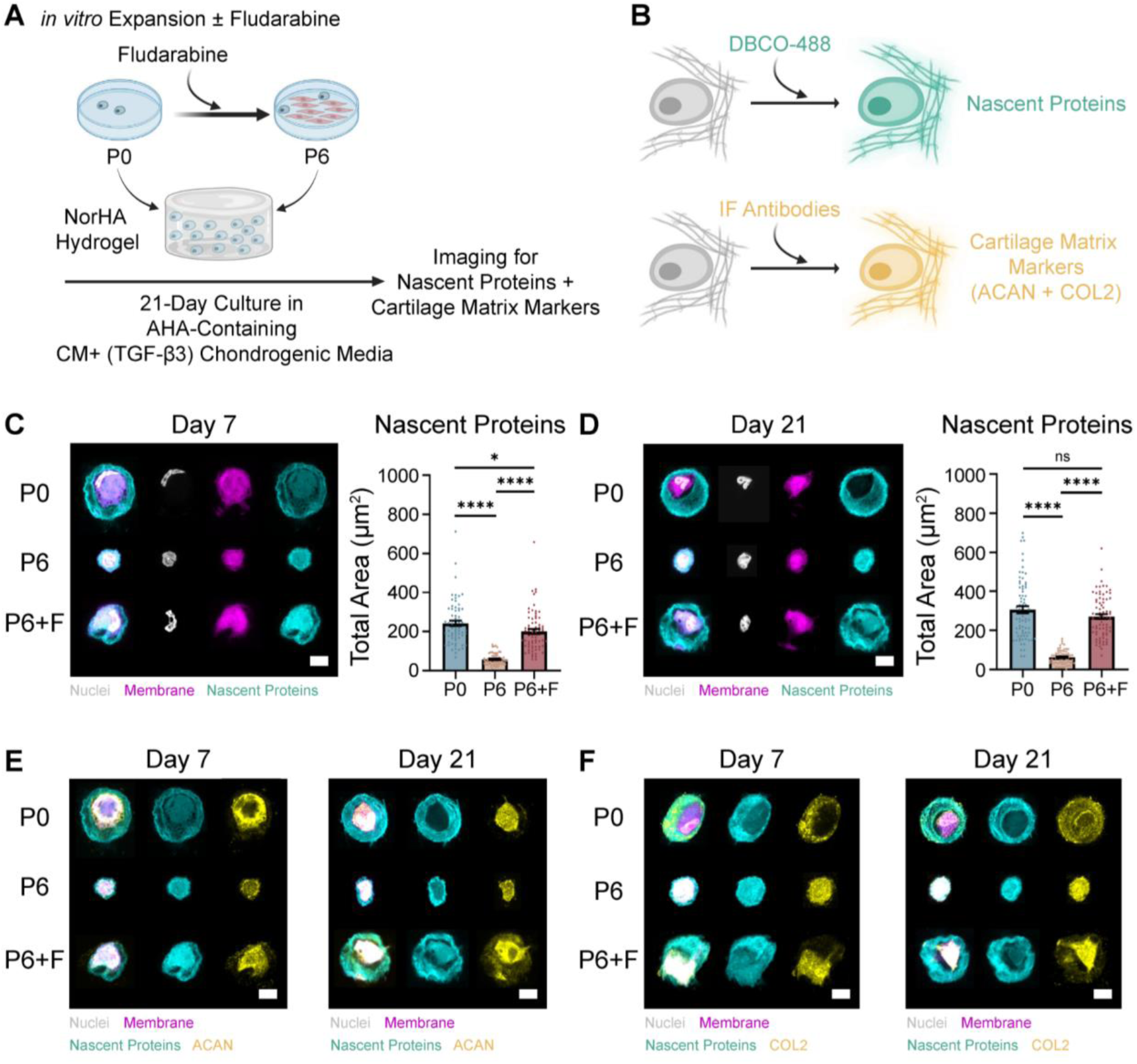
Fludarabine treatment during expansion enhances single-cell nascent protein synthesis and cartilage matrix marker deposition in NorHA-encapsulated chondrocytes. **(A)** Schematic of experimental design. Human chondrocytes were expanded *in vitro* to P6 with or without Fludarabine treatment, encapsulated in NorHA hydrogels, and cultured for 21 days in CM+ chondrogenic medium containing AHA, followed by **(B)** imaging of nascent proteins via click chemistry using DBCO-488, and cartilage matrix markers via immunofluorescence staining. **(C)** Representative images of nascent protein labeling and quantification of total nascent protein area at day 7 in P0, P6, and Fludarabine-treated (P6+F) chondrocytes. Data presented as mean ± SEM from one donor; n = 68 for P0, n = 65 for P6, n = 70 for P6+F, ****p < 0.0001, *p < 0.05 (one-way ANOVA with Tukey’s post hoc test). Scale bars = 10 µm. **(D)** Representative images of nascent protein labeling and quantification of total nascent protein area at day 21 in P0, P6, and P6+F chondrocytes. Data presented as mean ± SEM from one donor; n = 75 for P0, n = 66 for P6, n = 76 for P6+F, ****p < 0.0001, ns = not significant (one-way ANOVA with Tukey’s post hoc test). Scale bars = 10 µm. **(E)** Representative images of nascent proteins and ACAN staining at days 7 and 21 in P0, P6, and P6+F chondrocytes. Scale bars = 10 µm. Representative images of nascent proteins and COL2 staining at days 7 and 21 in P0, P6, and P6+F chondrocytes. Scale bars = 10 µm.

At day 7, P6 chondrocytes showed reduced nascent protein labeling relative to unexpanded P0 controls, consistent with impaired biosynthetic activity following expansion (**Fig. 4C**). In contrast, Fludarabine-treated P6 chondrocytes (P6+F) exhibited significantly greater nascent protein signal than untreated P6 cells, indicating partial recovery of protein synthesis capacity early in hydrogel culture (**Fig. 4C**). By day 21, the reduction in nascent protein synthesis in untreated P6 cells remained evident, whereas P6+F cells showed substantially increased nascent protein labeling, approaching P0 levels (**Fig. 4D**). In addition to increasing total nascent protein signal, Fludarabine treatment shifted the spatial distribution of nascent protein deposition toward a more P0-like distribution (**Fig. 4D**), suggesting improved biosynthetic activity and matrix assembly following expansion.

To further evaluate cartilage-specific matrix deposition, hydrogels were subsequently stained for aggrecan (ACAN) and type II collagen (COL II) following nascent protein imaging. At both day 7 and day 21, P6 chondrocytes showed weaker ACAN and COL2 staining than P0 controls, whereas P6+F cells displayed enhanced staining intensity for both markers (**Fig. 4E, F).** The recovery of cartilage matrix marker deposition in P6+F cells was especially evident by day 21, consistent with improved long-term maintenance of a cartilage-producing phenotype in 3D hydrogel culture (**Fig. 4E, F**).

Together, these findings demonstrate that Fludarabine treatment during expansion enhances nascent protein synthesis and cartilage matrix deposition at the single-cell level, supporting improved biosynthesis activity and matrix assembly capacity of expanded chondrocytes in 3D hydrogel environment.

### Fludarabine preserves native nanoscale chromatin architecture during *in vitro* expansion

Because stable cell identity is encoded not only in transcriptional programs but also in higher-order chromatin organization^32–34^, we next examined whether the transcriptional and biosynthetic preservation observed in Fludarabine-treated chondrocytes is accompanied by corresponding preservation of nanoscale chromatin spatial organization. To address this, we applied super-resolution stochastic optical reconstruction microscopy (STORM) and the O-SNAP (Objective Single-Molecule Nuclear Architecture Profiler) analysis pipeline^35^ to P0, P3, and P3+F chondrocytes. This approach enabled quantitative characterization of chromatin spatial organization at nanoscale resolution^35–38^. O-SNAP analysis extracted 144 spatial features encompassing nuclear morphology, global and radial chromatin organization, and multi-scale domain architecture^35^.

Initial visual inspection suggested reduced chromatin signal within the nuclear interior of P3 chondrocytes relative to P0 controls; however, Fludarabine-treated cells (P3+F) showed restoration of interior chromatin signal toward a P0-like pattern (**Fig. 5A**). To quantitatively characterize changes in histone-H3 (H3) spatial organization across conditions, we applied O-SNAP pipeline to STORM localization data. Feature selection using the Minimum Redundancy Maximum Relevance (MRMR) algorithm identified two features with the highest discriminative potential: Log_10_ (Border Curvature Mean), which reflects the average of the log-transformed distribution of the nucleus’s border curvature, and Log_10_ (Interior DBSCAN Cluster Radius Mean), which captures the average of the log-transformed distribution of the radius of all compact domains in a nucleus (**Fig. 5B–D**). Quantification of these features revealed that P3 chondrocytes exhibited significantly increased interior chromatin packing domain size and altered nuclear boundary curvature compared to P0 (**Fig. 5C, D**). Strikingly, P3+F chondrocytes showed no significant difference from P0 in nuclear boundary curvature (**Fig. 5C**), and a partial but significant recovery of interior packing domain size toward P0 levels (**Fig. 5D**). Together, these results indicate that Fludarabine treatment attenuates expansion-induced chromatin reorganization.

**Figure 5.**
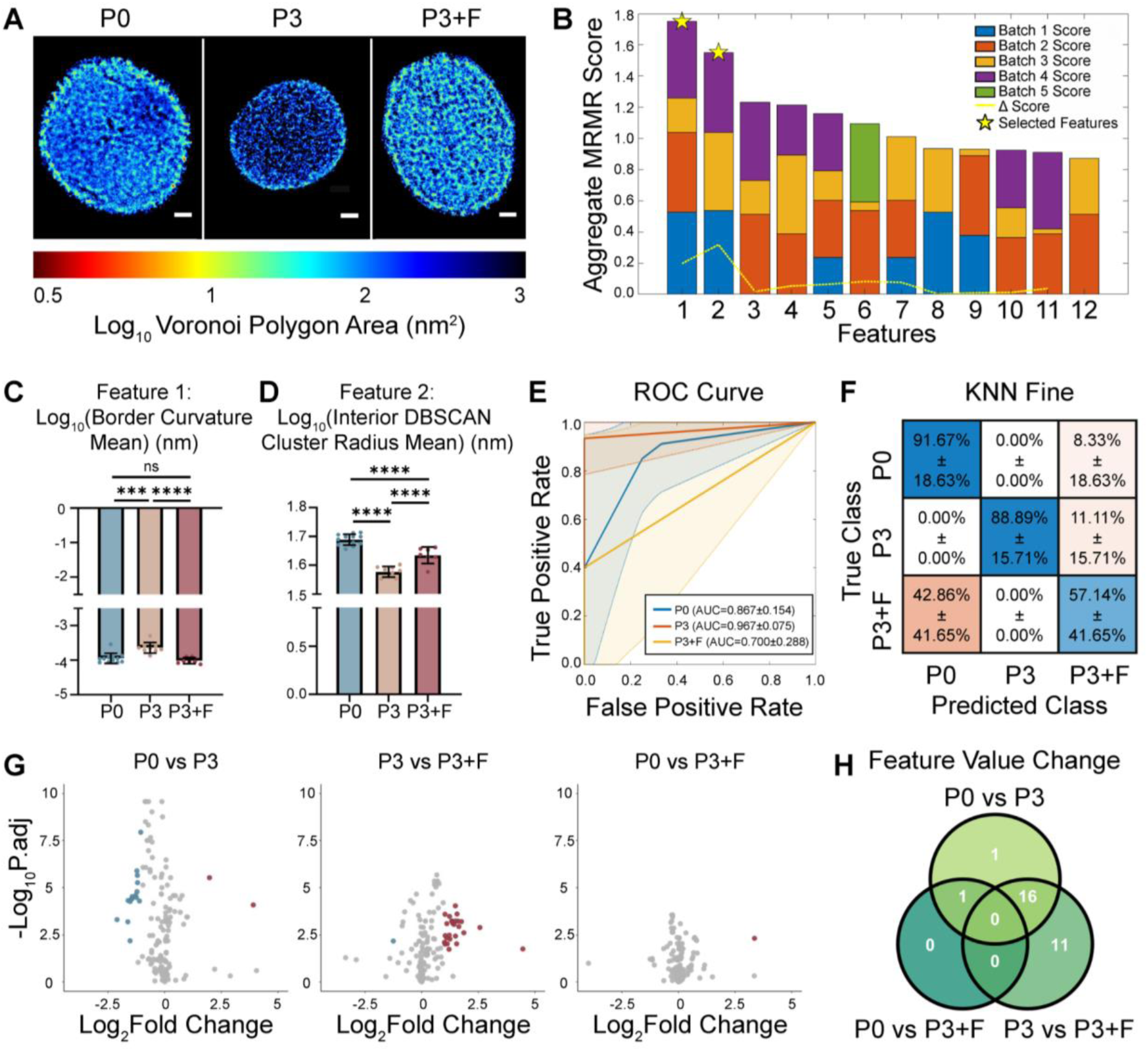
Fludarabine treatment preserves native nanoscale chromatin architecture in chondrocytes during *in vitro* expansion. **(A)** Representative Voronoi density map renderings of histone H3 STORM images of P0, P3, and P3+F chondrocytes. Scale bars = 2 µm. **(B)** Aggregated MRMR scores across five cross-validation folds for the top 12 O-SNAP features. Stacked bars indicate the contribution of each fold to the aggregate score. Two features were selected (stars): Feature 1 (Log_10_ Border Curvature Mean) and Feature 2 (Log_10_ Interior DBSCAN Cluster Radius Mean). **(C–D)** Quantification of **(C)** Log_10_ Border Curvature Mean and **(D)** Log_10_ Interior DBSCAN Cluster Radius Mean across P0 (n = 12 nuclei), P3 (n = 9 nuclei), and P3+F (n = 7 nuclei) chondrocytes. Data presented as mean ± SEM from one donor; ****p < 0.0001, ***p < 0.001, ns = not significant (one-way ANOVA with Tukey’s post hoc test). **(E)** ROC curves for a KNN fine classification model trained to discriminate P0, P3, and P3+F chondrocytes based on selected O-SNAP features across five cross-validation folds. Solid lines indicate average performance across training and test batches; shaded regions indicate ± 1 standard deviation. **(F)** Confusion matrix showing classification accuracy for the KNN fine model across the five folds. Values represent weighted mean ± standard deviation across training and test batches. **(F)** Volcano plots visualizing fold changes in O-SNAP-generated features of P0 vs P3, P3 vs P3+F, and P0 vs P3+F. Colored points indicate features with |Log_2_(FC)| > 1 and Benjamini-Hochberg adjusted p-value < 0.05. **(H)** Venn diagram showing the overlap of significantly differentially valued O-SNAP features across the three pairwise comparisons.

To further assess the extent to which Fludarabine restores a P0-like chromatin state, we performed machine learning-based classification using a K-nearest neighbor (KNN) model trained on the selected O-SNAP features. Many model architectures were trained using the selected feature, and the KNN model resulted in the best performance. The classifier distinguished P0 and P3 chondrocytes with high accuracy (**Fig. 5E**), confirming that these two groups occupy distinct chromatin states. In contrast, P3+F chondrocytes were classified with markedly lower confidence, and confusion matrix analysis revealed that 42.86% of P3+F nuclei were misclassified as P0, while none were misclassified as P3 (**Fig. 5F**). This pattern indicates that the classifier systematically conflates P3+F nuclei with the P0 chromatin state, reflecting substantial similarity in their chromatin architecture feature landscapes.

To characterize which specific chromatin features were altered across conditions and to what degree, we next performed pairwise volcano plot analyses of O-SNAP-derived features (**Fig. 5G**). Comparison of P0 versus P3 nuclei revealed numerous significantly differentially valued features between conditions, consistent with passage-dependent chromatin remodeling during *in vitro* expansion (**Fig. 5G**). In contrast, the P3 versus P3+F comparison showed the largest number of significantly altered features, indicating that Fludarabine treatment actively reshapes chromatin organization during expansion (**Fig. 5G**). Strikingly, comparison of P0 and P3+F nuclei revealed almost no significant feature differences, demonstrating that the chromatin architecture of P3+F chromatin closely resembles the native P0 state (**Fig. 5G**). Venn diagram analysis of differentially valued features further illustrated that while P0 vs P3 and P3 vs P3+F comparisons shared many differentially valued features, the P0 vs P3+F comparison contributed the fewest unique features (**Fig. 5H**).

Together, these findings demonstrate that STAT1 inhibition with Fludarabine preserves the nanoscale chromatin architecture of chondrocytes during *in vitro* expansion. Consistent with the transcriptional and biosynthetic evidence described above, these results suggest that Fludarabine-mediated STAT1 inhibition stabilizes the chromatin regulatory landscape underlying chondrocyte identity.

## 3. Discussion

Cartilage injuries and degenerative joint disease represent a major and growing clinical burden. Because articular cartilage has a limited intrinsic repair capacity, effective biological interventions for cartilage regeneration remain a critical unmet need. ACI and its matrix-assisted variant MACI have emerged as established cell-based strategies for cartilage repair, offering the potential for durable, biologically integrated tissue restoration^5,39,40^. A central challenge in ACI is that *in vitro* expansion, the very process required to generate clinically scalable chondrocyte numbers, simultaneously compromises the molecular and functional features that define chondrocyte phenotype. Decades of research have established that expanded chondrocytes undergo dedifferentiation, characterized by loss of cartilage-specific gene expression and matrix production capacity^7–9^. Although partial recovery of chondrogenic phenotype can be achieved through strategies such as 3D culture^8^, this redifferentiation remains incomplete. Consequently, expansion introduces a fundamental bottleneck for cell-based cartilage repair: although expansion is unavoidable, the resulting phenotype loss is not easily reversible. Instead, dedifferentiation appears to represent a progressive cellular transition, in which molecular and regulatory changes accumulate over time, gradually constraining the ability of expanded cells to reacquire a native chondrocyte phenotype.

Here, by profiling human articular chondrocytes across expansion using single-nucleus multiome sequencing (snMultiome-Seq), we show that dedifferentiation unfolds as a coordinated trajectory rather than a binary switch, and that this trajectory is accompanied by activation of specific regulatory programs. A key finding of this work is that progression along this dedifferentiation trajectory is associated with activation of transcription factors (TFs) that include STAT1. These data support a model in which expansion-associated dedifferentiation is not merely passive loss of cartilage markers, but an active state transition involving coordinated transcriptional and chromatin remodeling.

This trajectory-based view also helps explain why transcriptomic state alone does not always fully predict functional output. In our study, cells with similar transcriptomic profiles (P3 and P6) nevertheless differed in chromatin architecture, indicating that gene expression alone does not fully capture chondrocyte quality after expansion. This observation is particularly relevant in monolayer culture, where prolonged exposure to non-physiologic culture conditions can progressively reshape cell phenotype. Together, these findings suggest that expansion-associated phenotype loss reflects both molecular state transitions and culture history, and that preserving chondrocyte function may require intervention at the level of regulatory state transitions rather than downstream marker expression alone.

Among the transition-associated transcription factors identified along the dedifferentiation trajectory, STAT1 emerged as a particularly compelling candidate. Its combination of mechanistic relevance and pharmacologic tractability provided a rationale for targeting STAT1 during monolayer expansion to preserve chondrocyte phenotype. Pharmacologic inhibition of STAT1 signaling using Fludarabine preserved chondrocyte phenotype across multiple biological layers, including morphology, transcriptional programs, chromatin architecture, and matrix production. Importantly, these effects were not limited to recovery of a small number of canonical cartilage markers; instead, Fludarabine shifted broader regulatory programs related to cartilage differentiation and development toward a more native state, consistent with prior evidence that inflammatory signaling directly suppresses chondrogenic identity in part through STAT1 activation^41,42^. This indicates that the progressive activation of STAT1 during expansion may represent an active driver of chondrogenic suppression, and its inhibition by Fludarabine may relieve this transcriptional competition to permit restoration of the native chondrocyte state. To further resolve the mechanism of Fludarabine action, snMultiome profiling of Fludarabine-treated chondrocytes revealed that P3+F cells occupy an intermediate position along the dedifferentiation trajectory, with a subset of cells showing partial reversion toward the native P0 state. Notably, this recovery occurred preferentially at the chromatin level, with chromatin accessibility restored toward P0 to a substantially greater extent than RNA expression, suggesting that STAT1 inhibition acts primarily by stabilizing the epigenetic landscape underlying chondrocyte identity. GSEA further identified suppression of immune response-inhibiting signal transduction as the top transcriptional program enriched in Fludarabine-treated cells, directly consistent with attenuation of STAT1-driven inflammatory signaling during expansion. Consistently, at the single-cell functional level, FUNCAT measurements of nascent protein synthesis revealed that Fludarabine markedly increased anabolic activity in late-passage chondrocytes relative to untreated controls, with biosynthetic output approaching P0 levels by day 21 in 3D hydrogel culture. This indicates that suppression of STAT1-driven inflammatory programs stabilizes the epigenetic landscape and restores the functional matrix-producing capacity of expanded chondrocytes.

From a translational perspective, Fludarabine is an FDA-approved drug currently used in the treatment of hematologic malignancies, providing a well-characterized pharmacological and safety profile^43^. Importantly, in the context of ACI manufacturing, Fludarabine would be applied *ex vivo* during cell expansion rather than administered directly to patients, thereby effectively circumventing the systemic toxicity concerns associated with its chemotherapeutic use. This positions Fludarabine as an attractive drug-repurposing candidate for improving chondrocyte manufacturing quality and suggests a potential path toward clinical translation that leverages its existing regulatory history. More broadly, while the functional consequences of dedifferentiation or plasticity differ across cell types^44,45^, our findings support a general framework for studying and controlling expansion-associated state transitions in primary cell culture. In some tissues, dedifferentiation may facilitate repair; however, uncontrolled state shift during *in vitro* expansion often compromises phenotypic stability, functional potency, and product consistency^44,45^. The strategy presented here, integrating single nuclear multi-omic profiling with tissue-level and single-cell level readouts to identify pharmacologically targetable regulatory factors, may therefore be broadly applicable beyond chondrocytes to other primary cells in which expansion is necessary, but phenotype preservation is critical.

This work also raises several important opportunities and considerations for future studies. First, although inhibition of other TFs identified in this study did not preserve chondrocyte phenotype as effectively as STAT1 inhibition via Fludarabine (**Supplemental Fig. S4**), their therapeutic potential may not yet be fully defined. Further optimization of dose and treatment timing may reveal whether these pathways provide additional benefits, either as standalone or combination strategies. Second, although Fludarabine alone was sufficient to preserve the chondrogenic phenotype at transcriptomic, epigenetic, and matrix-production levels, it did not fully restore all genes altered during dedifferentiation (**Supplemental Fig. S5**). This suggests that additional regulatory pathways contribute to expansion-associated phenotype loss and supports future evaluation of combination strategies to further enhance chondrocyte phenotype preservation and improve ACI outcomes. For instance, although Fludarabine broadly shifted the transcriptional profile toward a P0-like state, a subset of genes remained unrestored, including structural matrix genes such as COL6A1, COL6A2, and MMP13, as well as residual inflammatory mediators, including SAA1, SAA2, NOS2, and IRF6 (**Supplemental Fig. S5**). This pattern suggests that STAT1-independent regulatory pathways contribute to specific aspects of expansion-associated phenotype loss, particularly those related to matrix structural gene expression and residual inflammatory signaling. Future studies exploring complementary regulatory axes may therefore further enhance chondrocyte phenotype preservation and improve ACI outcomes. Third, only male donors with the age range of 41–55 years were included in this study. Although sex differences and donor age were not central focuses of this work, both factors should be considered in future studies. This is particularly relevant given that ACI outcomes are generally more favorable in younger patients, suggesting that the phenotype-preserving effects of Fludarabine observed here may represent a conservative estimate of efficacy in the typical ACI patient population. Fourth, while the current study demonstrates phenotype preservation and recovery in controlled *in vitro* settings, translation to cartilage repair in a clinical setting will require validation in longer-term studies and *in vivo* models. In particular, it will be critical to determine whether preserved anabolic activity corresponds to improved tissue integrity and durable repair outcomes. Future studies in large animal models will therefore be essential to evaluate the *in vivo* efficacy and translational potential of this strategy. Finally, although dedifferentiation has been observed across species and age groups, species- and age-dependent differences in cartilage biology, cell plasticity, and repair capacity should be considered when selecting preclinical *in vivo* models and interpreting therapeutic efficacy.

In summary, our findings position chondrocyte dedifferentiation as a continuous, coordinated state transition during *in vitro* expansion and identify STAT1 activation as a prominent component of this trajectory. By showing that STAT1 inhibition via Fludarabine preserves chondrocyte phenotype at the transcriptional and chromatin levels and restores functional matrix-producing capacity, this work supports a phenotype-stabilization strategy to improve the quality of expanded chondrocytes. More broadly, these results suggest that targeted regulatory interventions during expansion can preserve the functional potential of therapeutic chondrocytes, helping to address a long-standing barrier to scalable and effective ACI.

## 4. Materials and Methods

### 4.1 Cell isolation, culture and expansion

Human knee articular cartilage tissues from seven adult donors (41-, 42-, 46-, 47-, 48-, 52-, and 55-year-old males) were obtained from the LifeLink Foundation. Primary human articular chondrocytes were isolated using enzymatic digestion with Liberase™ TM Research Grade (Roche, 5401119001) with gentle agitation for 18 h at 37 °C. Following digestion, the cell suspension was filtered through a 70 µm cell strainer, centrifuged at 400 x g for 6 min, and resuspended in basal growth medium consisting of high glucose Dulbecco’s Modified Eagle’s Medium (DMEM; Thermo Fisher Scientific, 11965118) supplemented with 10% fetal bovine serum (FBS; R&D Systems, S11150) and 1% penicillin–streptomycin (PS; Corning, 30-002-CI). For *in vitro* expansion, 500,000 chondrocytes were seeded onto 10 cm tissue culture dishes and maintained at 37 °C in a humidified incubator with 5% CO_2_. The culture medium was replaced every 2–3 days. Cells were passaged at approximately 70% confluency and expanded in monolayer culture to the designated passage for downstream experiments.

### 4.2 Pharmacological inhibition of STAT1 activation

STAT1 activity was inhibited using 10 μM Fludarabine (Selleck Chemicals, S1491). Chondrocytes were seeded and allowed to adhere for 24 h, after which Fludarabine was added in basal growth medium and incubated for 18 h. The following day, the medium was replaced with fresh basal growth medium without Fludarabine.

A second 18 h Fludarabine treatment was administered the next day using the same procedure. After the two 18 h treatments, cells were cultured to approximately 70% confluency and then passaged. This two-step treatment regimen was repeated at each passage for the Fludarabine-treated expansion groups. Depending on the downstream assay, Fludarabine-treated chondrocytes were expanded to either passage 3 (P3+F) or passage 6 (P6+F) using this treatment regimen.

### 4.3 Single-nuclei multiome sequencing (snMultiome-Seq)

Human chondrocytes from the indicated experimental groups (P0, P3, P6, and P3+F), pooled from four donors (41-, 46-, 52-, and 55-year-old males), were used for single-nucleus multiome profiling. Nuclei isolation, library preparation, and sequencing were performed by Genewiz (South Plainfield, NJ).

Nuclei were isolated following the 10x Genomics Demonstrated Protocol for Nuclei Isolation for Single Cell Multiome ATAC + Gene Expression Sequencing (CG000365, Rev C). Briefly, cryopreserved cells were thawed, lysed, washed, and centrifuged to recover nuclei, which were then resuspended in Nuclei Buffer for downstream library preparation. Nuclei concentration and quality were assessed prior to loading.

Single-nucleus ATAC + gene expression libraries were generated using the Chromium Next GEM Single Cell Multiome ATAC + Gene Expression platform (10x Genomics) according to the manufacturer’s instructions. The target nuclei recovery was 10,000 nuclei per sample. Libraries were sequenced on an Illumina platform using paired-end sequencing, with a target depth of 50,000 reads per cell total (25,000 reads per cell for the gene expression library and 25,000 reads per cell for the ATAC library).

### 4.4 snMultiome-Seq data processing and analysis

Raw sequencing data were processed using Cell Ranger ARC (10x Genomics, v2.0.2) with the human reference genome GRCh38 (refdata-cellranger-arc-GRCh38-2020-A-2.0.0). Individual libraries were processed using cellranger-arc count, and aggregated matrices were generated using cellranger-arc aggr with default depth normalization. For genotype-based demultiplexing, aligned BAM files from pooled snMultiome samples (Donors A–D) were processed using FreeBayes (v1.3.1)^46^ for variant calling, and donor identities were assigned using Souporcell (v2.4)^47^.

snMultiome data were analyzed using Seurat (v5.2.1)^48^ and Signac (v1.14.0)^49^. Gene annotations were obtained from Ensembl (release 98) via AnnotationHub (v3.10.1)^50^ and EnsDb.Hsapiens.v86 (v2.99.0). RNA and ATAC count matrices were imported using Read10X_h5, and Seurat objects were constructed with RNA and ATAC assays using CreateChromatinAssay. Peaks were restricted to standard chromosomes as defined by the hg38 reference genome. Quality control filtering was applied using RNA and ATAC metrics, including RNA features (200–7000), mitochondrial content (<15%), ATAC features (1000–30000), TSS enrichment (>1), and nucleosome signal (<2). Peaks were called using MACS2 (v2.2.9.1)^51^, and peaks overlapping blacklist regions or non-standard chromosomes were removed. A peak-by-cell matrix was generated using FeatureMatrix.

ATAC dimensionality reduction was performed using TF-IDF normalization and latent semantic indexing (LSI) via RunTFIDF and RunSVD, with UMAP visualization using LSI components (dims 2:30). Multimodal integration was performed using FindMultiModalNeighbors to construct a weighted shared nearest neighbor (wsnn) graph. For the additional P3F dataset, RNA was integrated using CCA, and ATAC was batch-corrected using Harmony (v1.2.3)^52^ prior to weighted nearest neighbor-based joint embedding and clustering.

Clustering was performed using the wsnn graph (FindClusters, resolution 0.05–0.08). For the additional P3F-integrated dataset, clustering and UMAP visualization were based on the CCA-integrated embedding at resolution 0.09. Cluster identities were manually assigned using canonical marker genes, including chondrogenic markers (COL2A1, ACAN, HAPLN1) and osteoarthritis-associated markers (COL10A1, COL1A1). Differential gene expression and chromatin accessibility analyses were performed using the wilcoxauc function in presto (v1.0.0)^53^. For transcription factor analysis, motif annotations were added using TFBSTools (v1.40.0)^54^, JASPAR2020 (v0.99.10)^55^, and BSgenome.Hsapiens.UCSC.hg38 (v1.4.5), and chromatin accessibility deviations were quantified using chromVAR (v1.24.0)^56^. Motif enrichment was assessed using FindMotifs, and significantly enriched motifs were identified based on adjusted p-values (Benjamini–Hochberg correction) and fold-enrichment thresholds.

Genome coordinate handling was performed using GenomeInfoDb (v1.38.8) and GenomicRanges (v1.54.1). Trajectory and pseudotime analyses were performed using a custom framework adapted from ArchR^57^, in which nuclei were ordered along a predefined trajectory (P0 → P3 → P6) by fitting a smooth spline through UMAP cluster centroids and projecting each nucleus onto the nearest point of the fitted curve. Dynamic changes in gene expression and chromatin accessibility across pseudotime were assessed using spline fitting and differential analysis across trajectory segments.

Integrated RNA–ATAC analyses were performed by correlating gene expression with chromatin accessibility and chromVAR motif activity across nuclei and clusters. Candidate transcription factors were identified based on concordant changes in motif accessibility and target gene expression, and ranked using differential accessibility, motif enrichment significance, and expression-based statistics. For gene regulatory network analysis, GRNs were inferred using Pando (v1.1.1)^58^. Data visualization was performed using ggplot2 (v3.5.1)^59^, ComplexHeatmap (v2.18.0)^60^, and Signac/Seurat plotting functions, including UMAP projections, feature plots, motif enrichment plots, and pseudotime-resolved heatmaps.

### 4.5 Immunoblotting

Chondrocytes were lysed in RIPA buffer (Thermo Fisher Scientific, 89900), and protein concentration was quantified using a Qubit Fluorometer (Thermo Fisher Scientific, Q33238). Equal amounts of protein (4 μg per sample) were loaded for electrophoresis.

Proteins were separated by SDS-PAGE on 4–20% polyacrylamide gels (Bio-Rad, 4561095) and transferred to 0.2 μm nitrocellulose membranes (Bio-Rad, 1704270) using a Trans-Blot Turbo system (Bio-Rad, 1704150). Membranes were blocked with 3% bovine serum albumin (BSA; Sigma-Aldrich, A1933) in TBST (Tris-buffered saline, Bio-Rad, 1706435, containing Tween-20, Thermo Scientific Chemicals, J20605.AP) for 1 h at room temperature and incubated overnight at 4 °C with primary antibodies against phospho-STAT1 (p-STAT1; 1:1000; Cell Signaling Technology, 9167) and GAPDH (1:3000; Thermo Fisher Scientific, 10494-1-AP). After primary antibody incubation, membranes were washed with TBST five times for 5 min each and incubated with HRP-conjugated secondary antibody (1:10000; Santa Cruz Biotechnology, sc-2768) for 1 h at room temperature, followed by five additional washes with TBST for 5 min each.

Protein bands were visualized using Clarity Western ECL Substrate (Bio-Rad, 1705060) and imaged using a ChemiDoc™ MP Imaging System (Bio-Rad, 12003154). Band intensities were quantified using ImageJ, and p-STAT1 signal was normalized to GAPDH.

### 4.6 Fluorescence imaging for cell morphology

For morphology imaging, chondrocytes were seeded, fixed, and imaged in Nunc™ Lab-Tek™ II 8-well chambered coverglass slides (Thermo Fisher, 155409PK). Cells were fixed with 4% paraformaldehyde (PFA; Thermo Fisher Scientific, J19943) for 30 min at room temperature, washed with PBS, permeabilized in PBS containing 0.05% Triton X-100 (Sigma-Aldrich, T8787), 320 mM sucrose (Sigma-Aldrich, S0389), and 6 mM magnesium chloride (Sigma-Aldrich, 208337) for 5 min at room temperature, and then blocked with 5% BSA for 1 h at room temperature.

For fluorescence labeling, cells were incubated with Alexa Fluor 488-conjugated phalloidin (1:200; Thermo Fisher Scientific, A12379) in PBS for 30 min at room temperature, washed with PBS, and nuclei were counterstained with ProLong Gold antifade mountant with DAPI (Invitrogen, P36935) immediately before imaging.

Images were acquired using a widefield Leica DM6000 microscope. Cell morphology was quantified using ImageJ.

### 4.7 Bulk RNA sequencing (RNA-Seq) and data analysis

Total RNA from chondrocytes was extracted using TRIzol reagent (Invitrogen, 15596026) and purified with the Direct-zol RNA Microprep Kit (Zymo Research, R2062) according to the manufacturer’s instructions. RNA integrity and concentration were assessed using a NanoDrop Microvolume UV-Vis Spectrophotometer (Thermo Fisher Scientific, ND-ONE-W), Qubit Fluorometer (Thermo Fisher Scientific, Q33238), and Agilent 2100 Bioanalyzer (Agilent Technologies, G2939B).

Library preparation and sequencing were performed by Genewiz (South Plainfield, NJ) as 150 bp paired-end reads on an Illumina platform. Raw sequence reads were trimmed to remove adapter sequences and low-quality bases using Trimmomatic (v0.36)^61^. Trimmed reads were aligned to the *Homo sapiens* reference genome (Ensembl) using STAR aligner (v2.5.2b)^62^. Gene-level read counts were quantified using featureCounts (Subread package v1.5.2)^63^.

Differential expression analysis was conducted using the DESeq2 package in R^64^. Differentially expressed genes (DEGs) were identified using a Benjamini-Hochberg adjusted p-value < 0.05 and an absolute log_2_ fold-change > 1.0. Global transcriptional changes were visualized using EnhancedVolcano^65^, while heatmaps illustrating expression patterns of genes of interest were generated using the pheatmap package in R^66^. Gene enrichment analysis was performed using clusterProfiler in R^67^, with enriched Gene Ontology (GO) biological processes identified using a false discovery rate (FDR) threshold of ≤0.05. Representative GO terms were visualized using ggplot2^59^.

### 4.8 Chondrocyte spheroids formation and evaluation

Spheroids were formed from P6 and P6+F chondrocytes at a density of 200,000 cells per spheroid using V-shaped wells and centrifugation at 310 x g for 5 min. Spheroids were cultured for 7 days in chondrogenic medium consisting of DMEM, 1% (vol/vol) Insulin-Transferrin-Selenium Premix (ITS Premix; Corning, CB-40350), 50 µg/mL L-proline (Sigma-Aldrich, P5607), 0.1 µM dexamethasone (Sigma-Aldrich, D2915), 0.9 mM sodium pyruvate (Sigma-Aldrich, P5280), 50 µg/mL 2-phospho-L-ascorbic acid (Sigma-Aldrich, 49752), 1.25 mg/mL BSA, 5.36 µg/mL linoleic acid (LA; Sigma-Aldrich, L1012), and 1% PS, with (CM+) or without (CM−) 10 ng/mL TGF-β3 (R&D Systems, 8420-B3). Medium was changed every other day.

Gene expression in spheroids was evaluated by RT-qPCR. Total RNA was extracted using TRIzol Reagent and purified with the Direct-zol RNA Microprep Kit. cDNA was synthesized using ProtoScript II First Strand cDNA Synthesis Kit (New England Biolabs, E6560S), and gene expression of chondrogenic markers was quantified using a QuantStudio 3 system (Thermo Fisher Scientific, A28136).

Remaining spheroids were fixed in 4% PFA for 1 h at room temperature, cryo-embedded, cryosectioned into 5 µm sections, and processed for Alcian blue (AB) and Picrosirius red (PSR) staining using standard histological procedures.

### 4.9 Metabolic labeling and fluorescent noncanonical amino acid tagging (FUNCAT) of nascent proteins and chondrogenic markers

NorHA was synthesized according to previously established protocol^31^. Briefly, sodium hyaluronate (HA; Lifecore Biomedical, ENG-00200) was dissolved in 2-(N-morpholino)ethanesulfonic acid (MES) buffer (pH 5.5; Sigma Aldrich, M3671) at 1% (w/v). 4-(4,6-Dimethoxy-1,3,5-triazin-2-yl)-4-methylmorpholinium chloride (DMTMM; TCI, D2919) was added to the HA solution, followed by dropwise addition of 5-norbornene-2-methylamine (Nor; TCI, N0907). The reaction was carried out for 24 h at room temperature. To isolate the polymer, saturated sodium chloride (Sigma Aldrich, S9888) was added, followed by precipitation with ethanol (Decon Laboratories, 2701). The polymer was collected by vacuum filtration, washed, resolubilized in deionized water, dialyzed for 3 days, and lyophilized. Norbornene modification was quantified by ^1^H NMR (600 MHz; Bruker, NEO400 spectrometer) of samples dissolved in D_2_O (7 mg/mL; Sigma Aldrich, 450510), using the ratio of vinyl to backbone proton integrals. Spectra were analyzed in MestReNova (v15.1.0).

5kPa NorHA hydrogels containing P0, P6 or P6+F chondrocytes were fabricated according to previously established protocol^28^. Briefly, a cell-hydrogel suspension was prepared at a density of 5 x 10^6^ cells/mL in PBS and hydrogel precursor solution. The suspension was photopolymerized using UV light (5 mW/cm^2^) for 3 min. Hydrogels were transferred to 24-well plates and cultured for 21 days in CM+ chondrogenic medium containing L-azidohomoalanine (AHA, Vector Labs, CCT-1066).

For nascent protein labeling, hydrogels were incubated with 30 μM Dibenzocyclooctyne-amine-488 (DBCO-488; Vector Labs, CCT-1278), CellMask Deep Red plasma membrane stain (Thermo Fisher Scientific, C10046), and Hoechst 33342 (Thermo Fisher Scientific, 62249) for 10 min at 37 °C and 5% CO_2_. Hydrogels were then washed 3 times with 5% BSA in PBS, fixed with 4% PFA for 30 min at room temperature, and washed 3 times with PBS.

For chondrogenic marker labeling, hydrogels previously imaged for nascent proteins were cryosectioned and subjected to immunofluorescence staining. Briefly, hydrogels were incubated in 30% sucrose (Thermo Fisher Scientific, S25590) overnight, snap-frozen the following day in Tissue-Tek Optimal Cutting Temperature Compound (O.C.T.; Electron Microscopy Sciences, 62550), and cryosectioned into 10 μm sections. Sections were blocked with 5% BSA for 30 min at room temperature and incubated overnight at 4 °C with primary antibodies against aggrecan (ACAN; 1:50; Abcam, ab3778) and type II collagen (COL II; 1:100; DSHB, II-II6B3). After primary antibody incubation, sections were washed with PBS 3 times for 10 min each and incubated with anti-mouse secondary antibody (1:200; Invitrogen, A11031) for 1 h at room temperature, followed by 3 additional PBS washes for 10 min each.

Z-stack images were acquired at 60x using a Nikon Eclipse Ti2-E confocal microscope. Total area for nascent proteins and chondrogenic markers was quantified in ImageJ by generating binary masks using Otsu thresholding, and the BoneJ plugin was used to calculate total area.

### 4.10 Super-resolution stochastic optical reconstruction microscopy (STORM) imaging and O-SNAP (Objective Single-Molecule Nuclear Architecture Profiler) chromatin architecture analysis

For STORM, chondrocytes were seeded, fixed, and imaged in Nunc™ Lab-Tek™ II 8-well chambered coverglass slides (Thermo Fisher, 155409PK). Cells were fixed with a methanol–ethanol (1:1) mixture for 6 min at −20 °C, followed by three washes with PBS. Blocking was performed with BlockAid solution (Thermo Fisher, B10710) for 1 h at room temperature to reduce nonspecific binding. Cells were incubated with mouse anti-histone H3 antibody (1:50; Active Motif, 39763) overnight at 4°C, washed with PBS, and then incubated with activator-reporter dye-paired secondary antibodies (Alexa Fluor 405-Alexa Fluor 647; Invitrogen, A30000 and A20006) for 1 h at room temperature, followed by another wash with PBS.

Imaging was performed on an ONI Nanoimager using repeated cycles of one activation frame (405 nm) followed by three imaging frames (647 nm). A fresh oxygen scavenger imaging buffer was prepared according to established protocols^38,68–70^, and contained 10 mM cysteamine (Sigma-Aldrich, 30070), 0.5 mg/mL glucose oxidase (Sigma-Aldrich, G2133), 40 mg/mL catalase (Sigma-Aldrich, C100), and 10% glucose (Thermo Fisher Scientific, A16828-36) in PBS.

Localization data from STORM imaging were analyzed using the O-SNAP (Objective Single-Molecule Nuclear Architecture Profiler) pipeline as previously described^35^. Nuclear boundaries were manually segmented in MATLAB, and 144 spatial features were extracted per nucleus, encompassing nuclear morphology, global and radial chromatin organization, and multi-scale domain architecture. Local chromatin compaction was quantified using Voronoi tessellation of fluorophore localizations. Chromatin packing domains were identified using DBSCAN clustering (ε = 20 nm, minimum neighbors = 3, minimum cluster size = 35 localizations). The nuclear area was partitioned into an interior region (inner 85%) and a peripheral region (outer 15%) for region-specific feature extraction

Pairwise comparisons of O-SNAP features across conditions were performed using two-sided t-tests with Benjamini-Hochberg correction for multiple comparisons. Features were considered significantly differentially valued at |log_2_FC| > 1 and adjusted p < 0.05. Feature selection was performed using the Minimum Redundancy Maximum Relevance (MRMR) algorithm across five-fold cross-validation, with the number of selected features determined by the knee-point of the MRMR score curve. Machine learning-based classification was performed using a K-nearest neighbor (KNN) model trained on PCA-transformed selected features, with model performance evaluated using ROC curves and confusion matrices across five cross-validation folds.

### 4.11 Statistical analysis

The number of biological replicates (*n*), statistical tests used, and *p*-values for each experiment are provided in the figure legends. Statistical analyses were performed using GraphPad Prism 10 (GraphPad Software, La Jolla, CA). For non-sequencing assays, data were evaluated for normality and outliers prior to parametric testing. Comparisons among groups were performed using one-way analysis of variance (ANOVA) followed by Tukey’s post hoc multiple-comparisons test. A *p*-value < 0.05 was considered statistically significant. For snMultiome-Seq and RNA-Seq analyses, significance was defined as a Benjamini–Hochberg adjusted *p*-value < 0.05 and an absolute log_2_ fold-change > 1.0.

## SUPPLEMENTARY MATERIAL

See the supplementary materials for additional figures and tables.

## ACKNOWLEDGMENTS

This research was supported by grants from the National Institutes of Health (R01 AR079224, P50 AR080581, R01 HL163168) and the NSF Science and Technology Center for Engineering Mechanobiology (CMMI-1578571). We would like to thank the LifeLink Foundation for providing human donor tissue used in this study, and we are grateful to the donors and their families for their generous contributions to research.

## AUTHOR DECLARATIONS

### Ethics approval

Ethics approval is not required.

## AUTHOR INFORMATION

These authors contributed equally: Ellen Y. Zhang and Sang Hyun Lee.

## Contributions

E.Y.Z., Y.C.L., Y.H., D.H.K., T.E.B., and J.J. performed biological experiments.

S.H.L. performed the snMultiome computational analysis.

E.Y.Z., Y.C.L., and H.H.K. performed data analysis.

E.Y.Z. and S.C.H. wrote the original manuscript.

All authors contributed to reviewing and editing the manuscript.

C.L., M.L., R.L.M., I.J., and S.C.H. supervised the work.

## Corresponding authors

Correspondence to Su Chin Heo and Inkyung Jung.

