## Supplemental Information for "Reprogramming Dedifferentiation Regulatory Networks Preserves Human Chondrocyte Phenotypes"

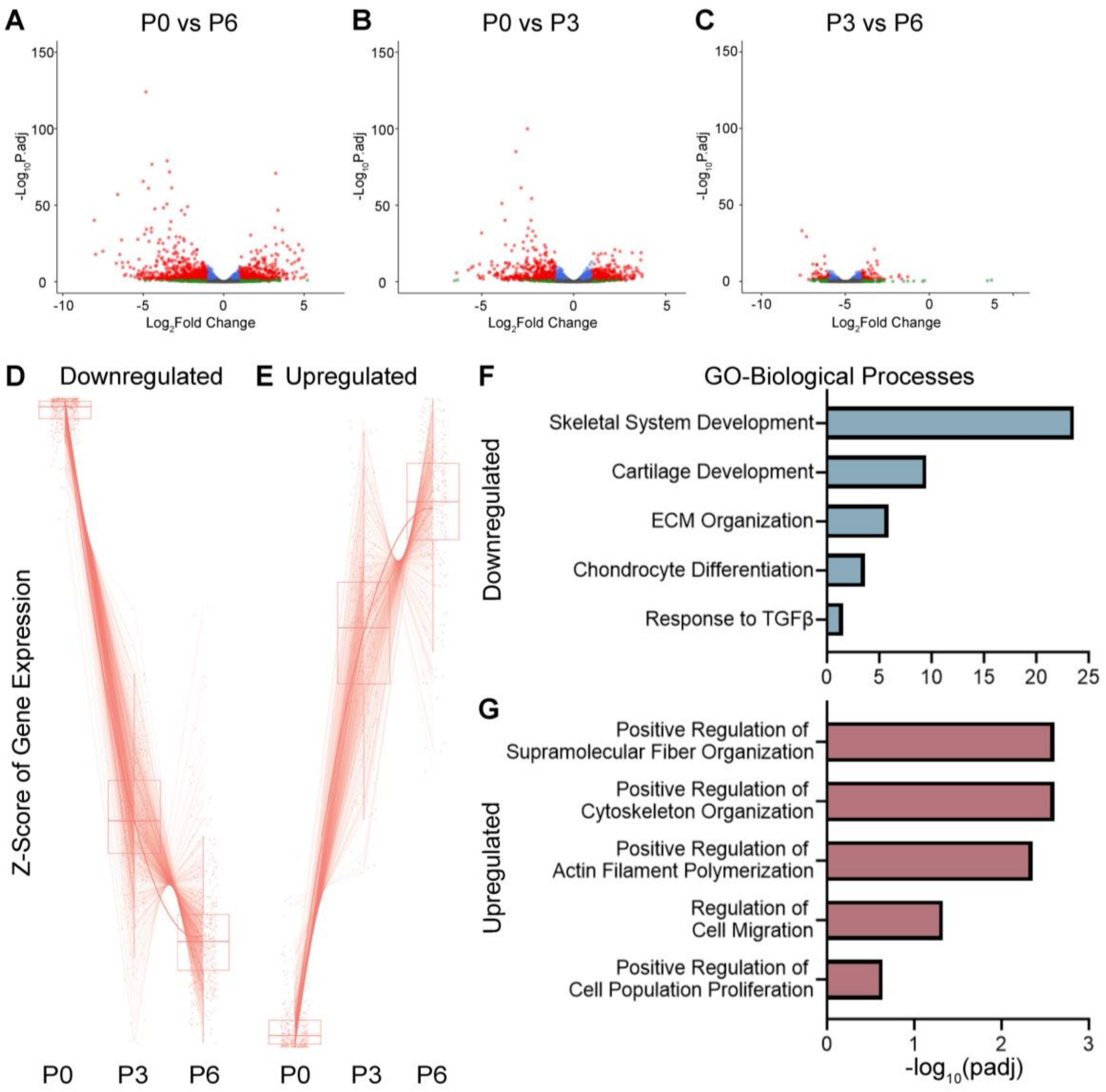


**Supplemental Figure S1. Bulk RNA-seq characterization of chondrocyte dedifferentiation across *in vitro* expansion. (A–C)** Volcano plots of differentially expressed genes (DEGs) comparing **(A)** P0 and P6, **(B)** P0 and P3, and **(C)** P3 and P6 human chondrocytes from four donors. Red points indicate DEGs (Benjamini-Hochberg adjusted $p < 0.05$, $|\log_2 FC| > 1$). **(D–E)** Trajectory plots of z-scored gene expression across P0, P3, and P6 for genes that are progressively **(D)** downregulated **or (E)** upregulated during expansion. Each line represents one gene. **(F–G)** Top Gene Ontology (GO) Biological Process terms enriched among **(F)** downregulated genes and **(G)** upregulated genes in the P0 vs P6 comparison.

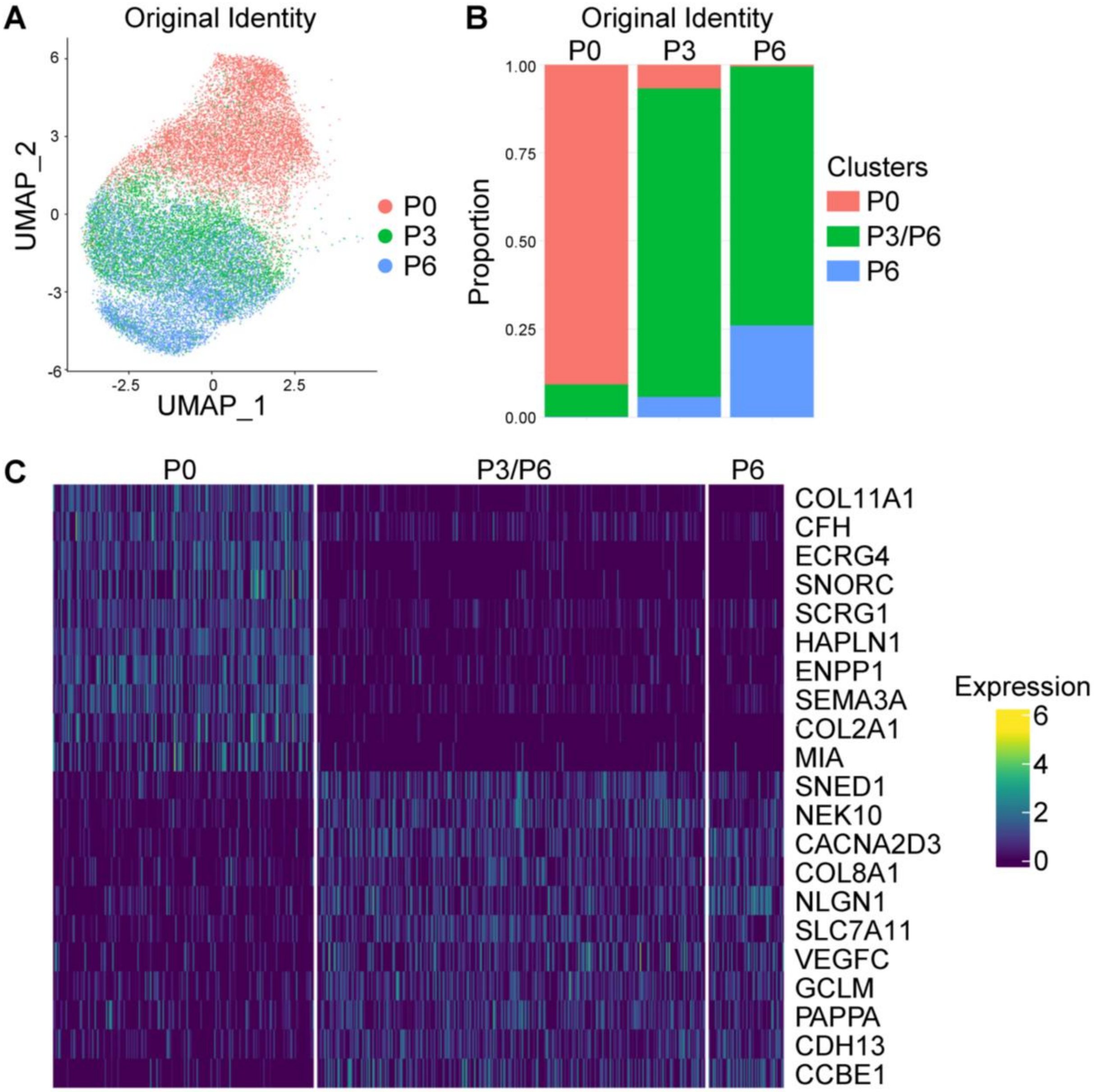


**Supplemental Figure S2. Integrated bimodal snMultiome analysis resolves P3 and P6 chondrocyte states that are indistinguishable by snRNA-seq alone**. **(A)** UMAP visualization of the snMultiome dataset colored by original passage identity (P0, P3, P6). **(B)** Stacked bar plots showing the proportion of nuclei from each passage assigned to each transcriptional cluster (P0-enriched, P3/P6-shared, and P6-enriched). **(C)** Heatmap of top differentially expressed marker genes across clusters.

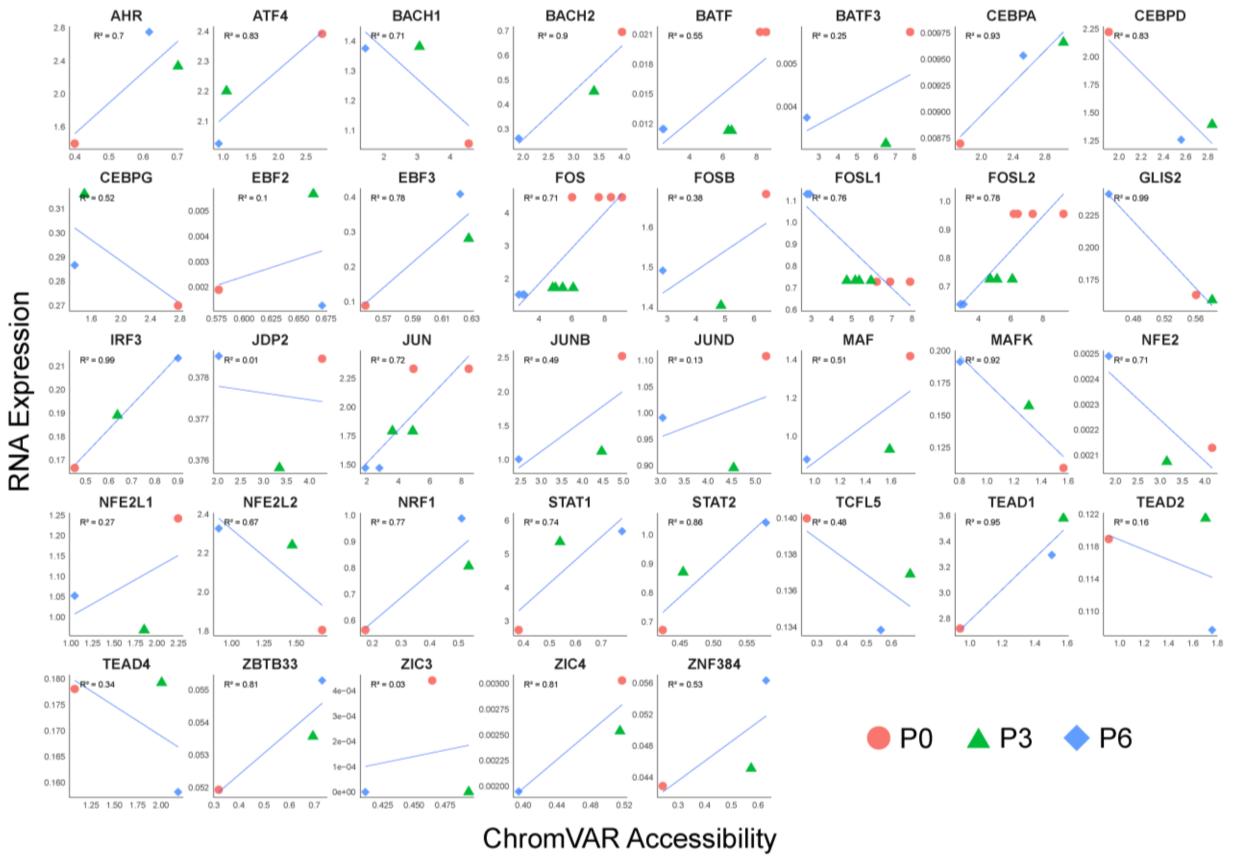


**Supplemental Figure S3. Concordance between transcription factor RNA expression and chromatin accessibility across chondrocyte passages.** Scatter plots showing the relationship between RNA expression (y-axis) and ChromVAR motif accessibility score (x-axis) for each candidate transcription factor across P0, P3, and P6. A linear regression line is fit for each TF with the corresponding $R^2$ value indicated.

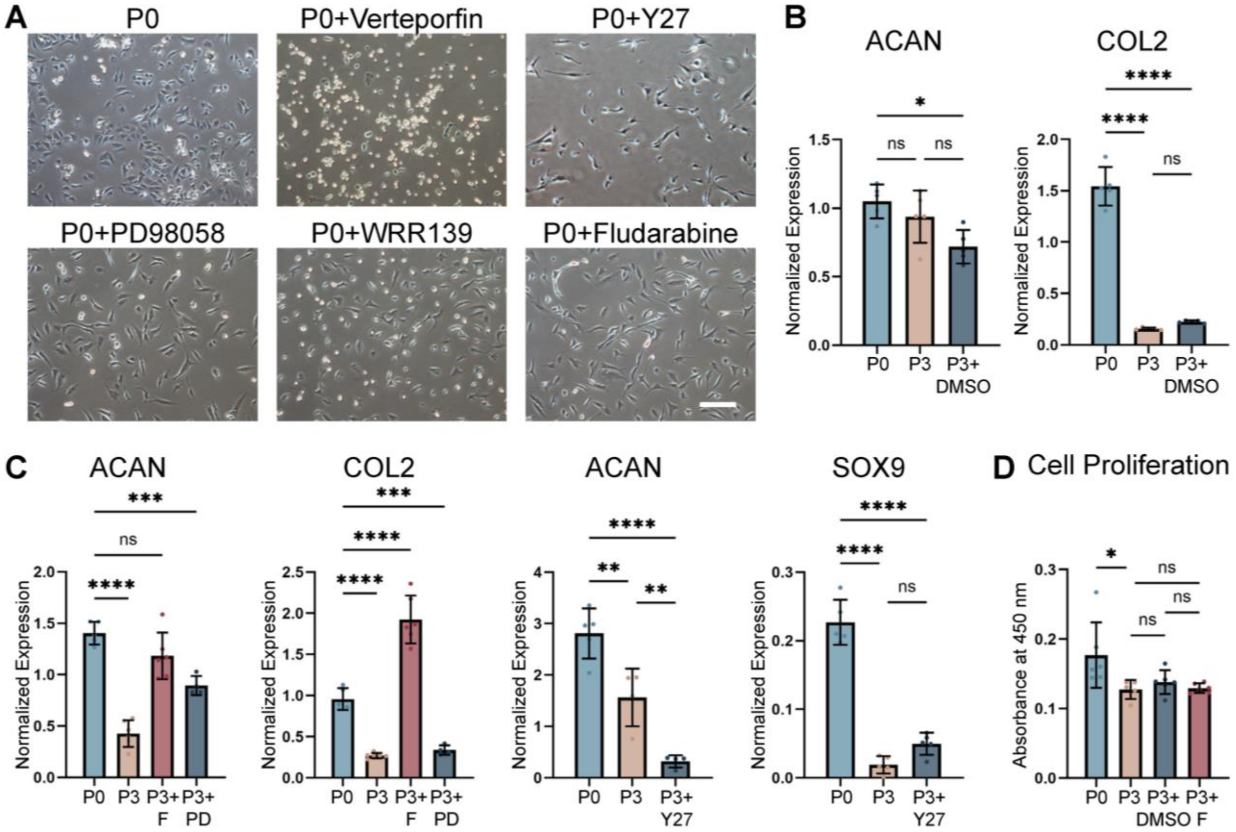


**Supplemental Figure S4. Pharmacological inhibitor screen for candidate transcription factors identified by snMultiome analysis. (A)** Representative brightfield images of untreated P0 chondrocytes and P0 chondrocytes treated overnight with Verteporfin, Y-27632, PD98058, WRR139, or Fludarabine. Scale bar = 200 µm. **(B)** RT-qPCR analysis of ACAN and COL2 expression in P0, P3, and P3 treated with DMSO vehicle control (P3+DMSO). Data normalized to GAPDH and presented as mean ± SEM from one donor; n = 5, ****p < 0.0001, *p < 0.05, ns = not significant (one-way ANOVA with Tukey's post hoc test). **(C)** RT-qPCR analysis of ACAN and COL2 expression in P0, P3, P3+Fludarabine (P3+F), and P3+PD98058 (P3+PD), and ACAN and SOX9 expression in P0, P3, and P3+Y-27632 (P3+Y27). Data normalized to GAPDH and presented as mean ± SEM from one donor; n = 5, ****p < 0.0001, ***p < 0.001, **p < 0.01, *p < 0.05, ns = not significant (one-way ANOVA with Tukey's post hoc test). **(D)** Cell proliferation assessed by CCK-8 assay in P0, P3, P3+DMSO, and P3+Fludarabine. Data presented as mean ± SEM from one donor; *p < 0.05, ns = not significant (one-way ANOVA with Tukey's post hoc test).

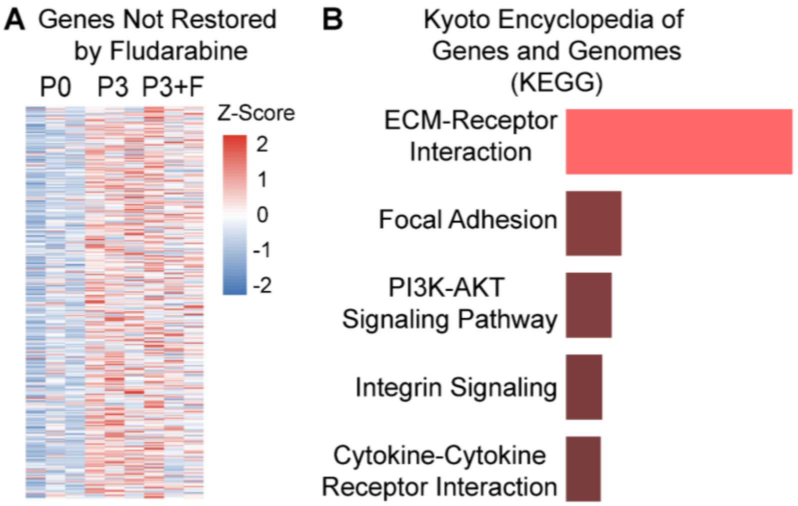


**C** Selected Genes Not Restored by Fludarabine by Functional Category

| Immune/ Inflammation | Focal Adhesion/ Mechanosensing |
|---|---|
| SAA1 | ITGA9 |
| SAA2 | RAPH1 |
| NOS2 | TIAM2 |
| IRF6 | ARHGAP5 |
| TRIM22 | CDK5 |
| IL13RA1 | FGD5 |
| TSLP | SDC1 |
| LAPTM5 | FAT4 |
| STAB1 | ADGRG2 |
| ACKR4 | ADGRG4 |
| BCL6 | ADGRG6 |
| ETS1 | GPC6 |
| TGFB2 | LMCD1 |
| CYSLTR1 | PLXDC2 |

**Supplemental Figure S5. Characterization of genes not restored by Fludarabine treatment.** **(A)** Heatmap showing z-scored expression of the 178 genes that remained differentially expressed between P3+F and P0 chondrocytes despite Fludarabine treatment, across P0, P3, and P3+F chondrocytes from three donors. **(B)** KEGG pathway enrichment analysis of Fludarabine-refractory genes. **(C)** Selected Fludarabine-refractory genes grouped by functional category.

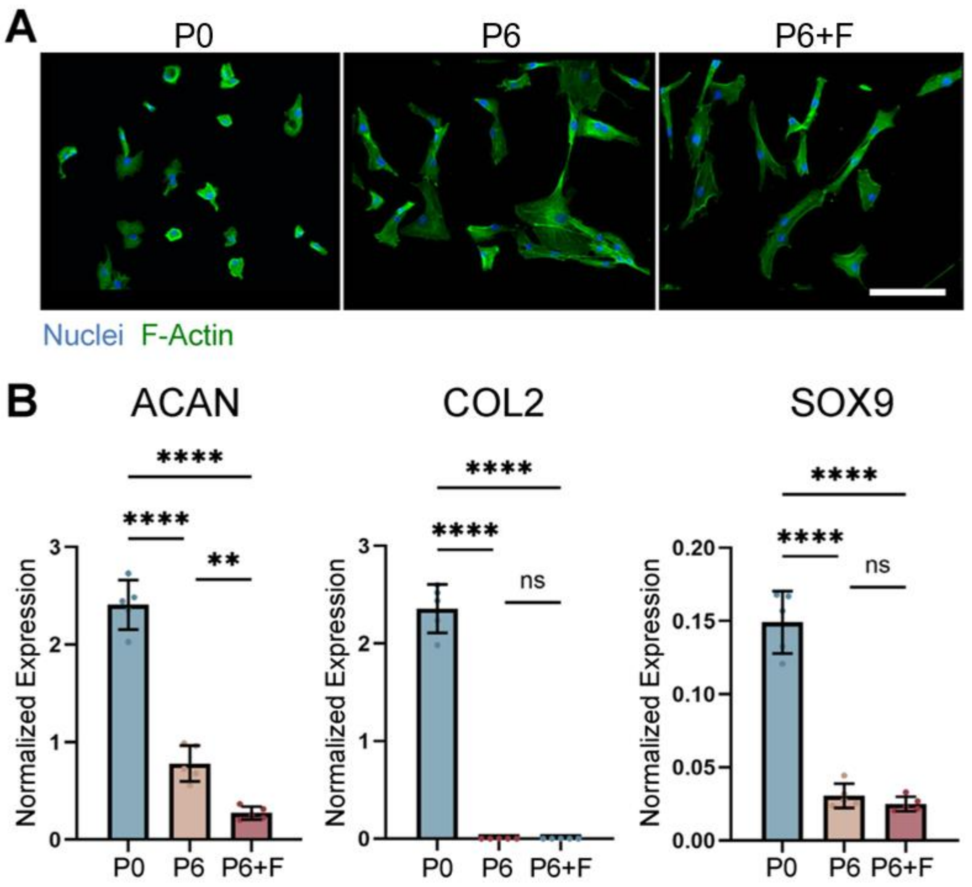


**Supplemental Figure S6. Late-stage Fludarabine treatment did not restore chondrocyte phenotype in already-dedifferentiated P6 chondrocytes. (A)** Representative fluorescence images of P0, P6, and P6+F chondrocytes stained for nuclei and F-actin. Scale bar = 100 μm. **(B)** RT-qPCR analysis of ACAN, COL2, and SOX9 expression in P0, P6, and P6+F chondrocytes. Data normalized to GAPDH and presented as mean ± SEM from one donor; n = 5, ****p < 0.0001, **p < 0.01, ns = not significant (one-way ANOVA with Tukey's post hoc test).

**Supplemental Table S1. Summary of pharmacological inhibitor screen targeting candidate transcription factors.** Candidate transcription factors identified from snMultiome analysis were assessed by pharmacological inhibition during chondrocyte *in vitro* expansion. For each target, the inhibitor used, dose, treatment schedule, and key observations are summarized.

| Targeted TF | Inhibitor | Dose & Treatment Time | Observations | Dose & Treatment Time References |
|---|---|---|---|---|
| **TEAD1** | Verteporfin | 10 μM; Overnight x 2 consecutive days | Cytotoxic under tested conditions. Initial treatment tolerated; however, cells detached and rounded upon second treatment. Upstream inhibition of ROCK with Y-27632 (see below) further corroborates that TEAD1 transcriptional activity is not required for chondrogenic phenotype maintenance during expansion. | 1 |
| | Y-27632 | 10 μM; 1 hr x 3 consecutive days | Well tolerated but did not preserve chondrogenic gene expression. ROCK inhibition suppresses YAP/TAZ nuclear translocation and thus TEAD1 transcriptional activity, providing independent upstream validation that this axis does not drive expansion-associated dedifferentiation (See Fig. S4). | 2 |
| **STAT2** | Not Applicable | | Not independently assessed; STAT2 signaling is expected to be suppressed downstream of STAT1 inhibition via Fludarabine. | Not Applicable |
| **NRF1** | WRR139 | 5 μM; Overnight x 2 consecutive days | Cytotoxic under tested conditions; cell death was observed after first treatment. Phenotypic assessment therefore not possible at this dose and treatment schedule. | 3 |
| **STAT1** | Fludarabine | 10 μM; Overnight x 2 consecutive days | Preserved chondrogenic gene expression and was therefore prioritized for further characterization (See main text and Fig. S4). | 4 |
| **AHR** | PD98058 | 10 μM; Overnight x 2 consecutive days | Well tolerated but produced partial and inconsistent effects on chondrogenic gene expression. Expression of select chondrogenic genes was modestly increased relative to untreated P3 controls but did not reach P0 levels, while other chondrogenic markers showed no improvement. Overall effects were weaker and less consistent than those observed with Fludarabine treatment (See Fig. S4). | 5 |

**Supplemental Table S2. RT-qPCR primer sequences.** Forward and reverse primer sequences used for RT-qPCR quantification of chondrogenic marker genes (ACAN, COL2A1, SOX9) and the housekeeping reference gene GAPDH. Expression values were quantified using the standard curve method and normalized to GAPDH. Data presented as normalized expression (target gene quantity / GAPDH quantity).

| Gene Name | Forward Sequence | Reverse Sequence |
|---|---|---|
| ACAN | TGAGGAGGGCTGGAACAAGTACC | GGAGGTGGTAATTGCAGGGAACA |
| SOX9 | GTACCCGCACTTGCACAAC | TCGCTCTCGTTCAGAAGTCTC |
| GAPDH | AACAGCGACACCCACTCCTC | CATACCAGGAAATGAGCTTGACAA |
| COL2A1 | GGCAATAGCAGGTTCACGTACA | CGATAACAGTCTTGCCCCACTT |
